# Simultaneous aerosol and intra-muscular immunization with influenza vaccine induces powerful protective local T cell and systemic Ab immune responses in pigs

**DOI:** 10.1101/2020.09.10.291625

**Authors:** Veronica Martini, Basu Paudyal, Tiphany Chrun, Adam McNee, Matthew Edmans, Emmanuel Maze, Beckie Clark, Alejandro Nunez, Garry Dolton, Andrew Sewell, Peter Beverley, Ronan MacLoughlin, Alain Townsend, Elma Tchilian

**Affiliations:** The Pirbright Institute, Pirbright GU24 0NF, UK; Animal and Plant Health Agency-Weybridge, New Haw, Addlestone KT15 3NB, UK; Division of Infection and Immunity, Cardiff University School of Medicine, Cardiff, Wales, UK; National Heart and Lung Institute, St Mary’s Campus, Imperial College, London W2 1PG, UK; Aerogen Ltd, IDA Business Park, Dangan, Galway, Ireland; Weatherall Institute of Molecular Medicine, University of Oxford, Oxford OX3 9DS, UK

## Abstract

A vaccine providing both powerful antibody and cross-reactive T cell immune responses against influenza viruses would be beneficial for both humans and pigs. Here we evaluated intramuscular (IM), aerosol (Aer) and simultaneous immunization (SIM) by both routes in pigs, using the single cycle candidate influenza vaccine S-FLU. After prime and boost immunization pigs were challenged with H1N1pdm09 virus. IM immunized pigs generated high titer of neutralizing antibodies but poor T cell responses, while Aer induced powerful respiratory tract T cell responses, but a low titer of antibodies. SIM pigs combined high antibody titers and strong local T cell responses. SIM pigs showed the most complete suppression of virus shedding and the greatest improvement in pathology. We conclude that SIM regimes for immunization against respiratory pathogens warrant further study.

## Introduction

Immunization against infectious diseases has been practised for several centuries but identifying the best method of administering a vaccine is still often a matter of empirical experimentation. Three major considerations should make rational immunization easier. The first is the importance of pathogen associated molecular patterns, which are essential for triggering an immune response. The second that the site of immunization programmes lymphocytes to return to it. The third that local immune responses are critical for protection against mucosal infection and that many lymphocytes reside in non-lymphoid tissues and provide tissue resident memory. Immunization at the site of infection offers the advantage that an immune response is generated at the site of entry of the pathogen and should provide immediate protection.

Immunization of the respiratory tract has been demonstrated to be highly effective against influenza and cold adapted live attenuated influenza vaccine (LAIV) has efficacy rates of 75-80% in children and additionally gives some cross-reactive protection against antigenically distinct strains. However, LAIV is not so effective in adults or the elderly. In contrast, the traditional intra-muscular inactivated seasonal human influenza vaccine provides 10-60% efficacy and induces strain-specific immunity by generation of subtype specific antibody, so that repeated annual vaccination to match new influenza variants is required^1-3^. Therefore, there is an urgent need for new immunization strategies for influenza that provide broad and long-lasting protection.

One such strategy, which has been explored against tuberculosis (Tb), is to combine the advantages of local and systemic immunization. Parenteral BCG priming followed by intranasal boosting with an Adenovirus vectored vaccine expressing antigen 85A (Ad85A) markedly enhanced protection in mice^4^. We have shown that simultaneous systemic and respiratory immunization (SIM) with BCG in mice or BCG/BCG and BCG/Ad85A in cattle enhanced protection against Tb challenge^5,6^. Uddback et al have used this strategy with an Adeno vector expressing influenza nucleoprotein (NP) and shown greatly improved and durable protection against heterosubtypic influenza challenge in mice^7,8^. These data prompted us to test SIM in the pig model using the candidate broadly protective signal minus influenza vaccine S-FLU. S-FLU is a pseudotyped influenza virus, lacking the HA signal sequence and therefore limited to a single cycle of replication. S-FLU induces a strong cross-reactive T cell response, but a minimal humoral response to hemagglutinin when administered mucosally^9,10^. We have shown that aerosol delivery of S-FLU reduces lung viral load when partially matched to the challenge virus, correlating with a local lung T cell immune response^11^. When S-FLU was completely mismatched to the challenge virus, pathology but not viral load, was reduced. This suggests that, in the absence of an antibody response, lung T cell immunity can reduce disease severity^12^. By contrast, the same S-FLU preparation induced sterile immunity to the matched challenge virus and reduced replication and aerosol transmission to naïve recipients following mismatched viral challenge in ferrets^12^. The pig is a more relevant large animal model because it is a natural host for influenza viruses and has very similar respiratory anatomy to humans^13,14^. Pigs and humans are infected by the same subtypes of influenza A viruses and are integrally connected in the ecology of influenza.

Here we evaluated the efficacy of SIM with S-FLU against H1N1pdm09 challenge using inbred Babraham pigs, allowing a more refined analysis of the specificity of the immune responses using MHC class I tetramers to previously defined immunodominant NP epitopes^15^.

## Results

### Virus load and lung pathology

To evaluate the efficacy of simultaneous pulmonary and systemic immunization, groups of 6 inbred Babraham pigs were immunized with S-FLU expressing NA and coated in the HA from H1N1pmd09 intra-muscularly (IM) or by aerosol (Aer) alone or simultaneously by aerosol and intra-muscularly (SIM). The SIM group received the same total dose as the IM or Aer groups, but split between the two sites. Untreated pigs were used as controls. The animals were boosted 3 weeks later and, after a further 3 weeks, challenged with H1N1pdm09 virus and culled 4 days after the challenge (**Fig. 1a**). Two pigs were culled before the end of the experiment because of underlying heart conditions, unrelated to the study, leaving 5 animals in the IM and control groups. Virus load was assessed in nasal swabs and broncho-alveolar lavage (BAL). The SIM pigs showed the greatest reduction of virus shedding in the nasal swabs at all time points except for the third day post challenge (DPC) (**Fig. 1b**). In the IM group, two individuals shed virus consistently after challenge but a significant reduction in viral load was achieved on 1 DPC. Aerosol immunization did not decrease virus shedding although 2 pigs did not shed at 4 DPC (**Fig. 1b**). Overall IM and SIM significantly reduced the viral load in the nasal swabs over time, with an average area under the viral load/time curve (AUC) of 3.46 and 2.23 respectively compared to 9.53 and 11.46 of the Aer and control group (**Fig. 1c**). No virus was detected in BAL at 4 DPC in any of the immunized groups (**Fig. 1d**).

**Figure 1.**
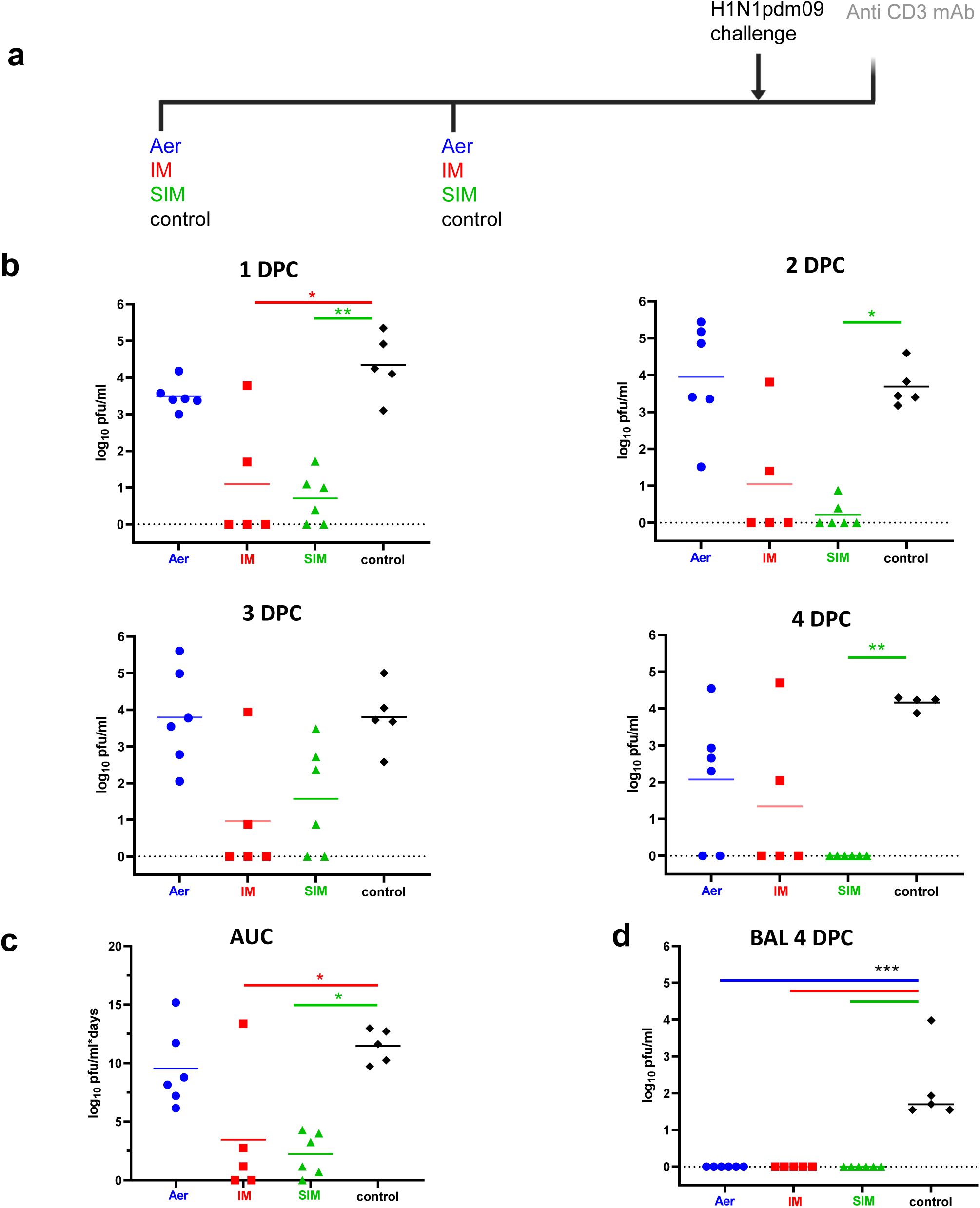
Experimental design and viral load in nasal swabs and BAL. **(a)** Babraham pigs were immunized with S-FLU by aerosol (Aer), intramuscularly (IM) or simultaneously by Aer and IM (SIM) and boosted 3 weeks apart. Control animals were left untreated. All animals were challenged with H1N1pdm09 virus 3 weeks after the boost. Swabs were taken daily post challenge and all pigs were culled 4 days post challenge (4 DPC). Half of the pigs were infused intravenously with anti-porcine CD3 mAb 10 minutes prior to sacrifice. (**b)** Virus titre in nasal swabs measured by plaque assay at 1, 2, 3 and 4 DPC. **(c)** Area under the curve (AUC) of viral titre in the nasal swabs over time. **(d)** Viral titre in the broncho-alveolar lavage (BAL) 4 DPC. The data represents the average of 2 separate assays, each data points indicates an individual animal and the horizontal line the mean of the group. Data were analysed using the Kruskal-Wallis test. Asterisks indicate significant difference from the control group *p<0.05, **p<0.01 *** p<0.001.

The unimmunized animals showed typical gross pathology changes in the lungs with multifocal areas of consolidation in the cranial and medial lobes (**Fig. 2**). A significant reduction in the extension and severity of the gross changes was observed in the IM and SIM groups (p=0.02 and p=0.005 respectively compared to controls), with a trend towards improved pathology in the aerosol group that did not reach statistical significance (**Fig. 2b**). A characteristic bronchiointerstititial pneumonia, with bronchiolitis, alveolar exudation and lymphohistiocytic infiltration in the alveolar septa and peribronchial and perivascular areas, was present in the unimmunized animals. A reduction in the severity of these changes was observed in the immunized groups (**Fig. 2a**). Labelling of influenza A nucleoprotein (NP) by immunohistochemistry (NP-IHC) was seen in only one animal in the IM and two animals in the SIM groups (p=0.02 and p=0.03 respectively), whereas most non-immunized pigs displayed abundant labelling. NP-IHC was reduced in the Aer group although this was not significant (p=0.55). Despite a reduction in gross lesions score and number of virus infected cells, no significant difference was found when histopathology and NP-IHC were combined in all immunized groups (Iowa score) (**Fig. 2b**).

**Figure 2.**
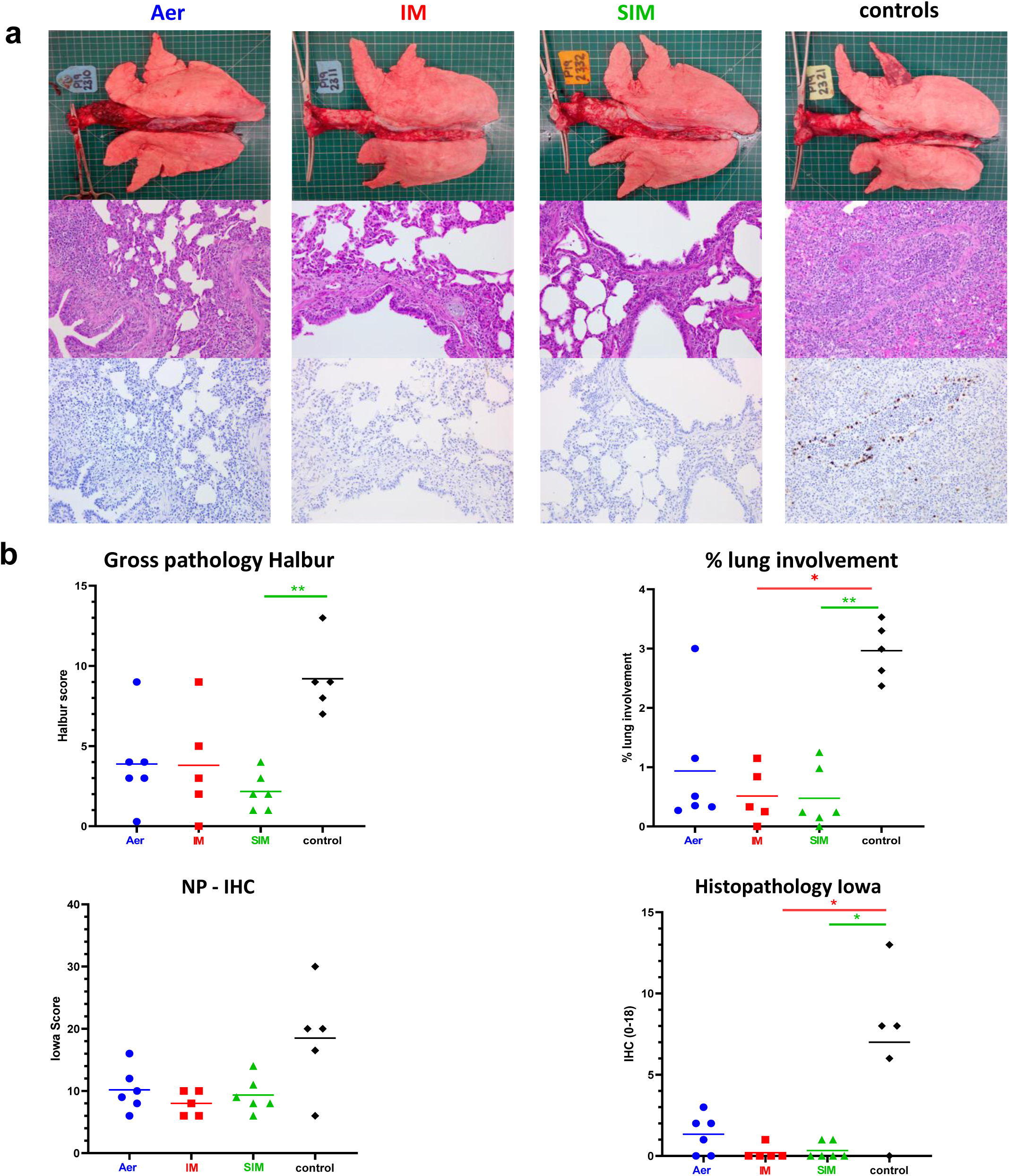
Lung pathology. Pigs were immunized with S-FLU by aerosol (Aer), intramuscularly (IM) or simultaneously by Aer and IM (SIM) while control pigs were untreated. Three weeks post boost pigs were challenged with H1N1pdm09. The animals were euthanized at 4 DPC and lungs scored for appearance of gross and histopathological lesions. Representative gross pathology, histopathology (H&E staining; 100x) and immunohistochemical NP staining (200x) for each group are shown **(a)**. The gross and histopathological scores for each individual in a group and the group means are shown (**b**), including the percentage of lung surface with lesions, the lesion scores and the histopathological scores (“Iowa” includes the NP staining). Pathology scores were analysed using one-way non-parametric ANOVA with the Kruskal-Wallis test. Asterisks denote significant differences *p<0.05 and **p<0.01 compared to control.

These results indicate that IM and SIM immunization significantly reduced nasal virus shedding and pathology, with SIM being more effective in virus clearance and gross lung pathology reduction. Aerosol immunization did not significantly reduce nasal virus load or pathology. All immunizations regimes eliminated virus in the BAL.

### Antibody and B cell responses

The serum neutralizing titer against H1N1pdm09 in the IM and SIM groups increased after the boost to a peak at of 4,096 (50% inhibition titre) and 1,812 respectively at 14 days post boost (DPB) and declined by 22 DPB and after the challenge (**Fig. 3a**). The Aer group had a much lower peak serum neutralizing titer of 54 at 14 DPB (**Fig. 3a**). Neuraminidase (NA) inhibition activity was assessed by enzyme-linked lectin assay (ELLA) at 4 DPC in the serum. IM immunized animals showed the highest inhibition titer (1,408) followed by SIM (453) and Aer (14.2) (**Fig. 3a**).

**Figure 3.**
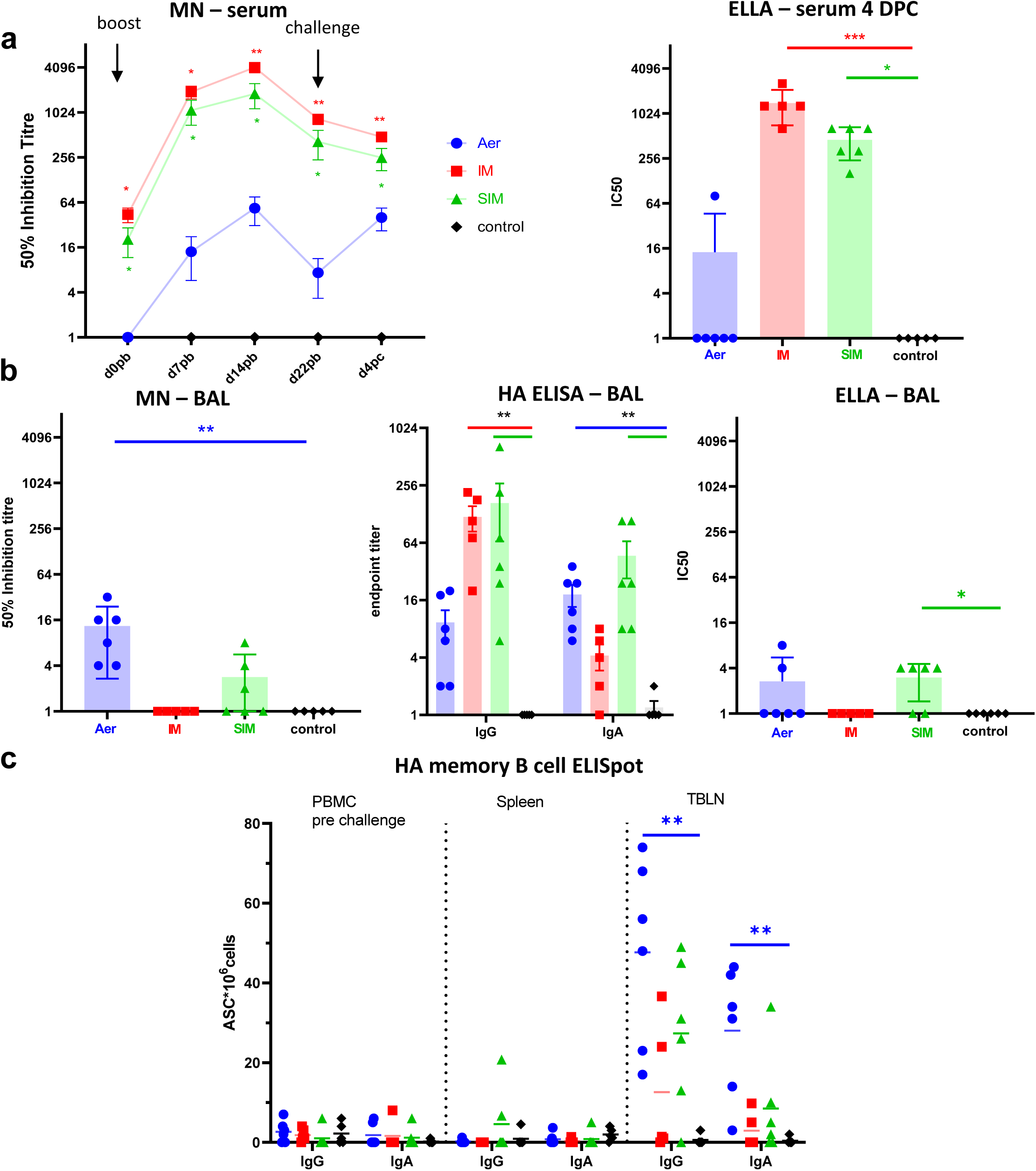
Systemic and local antibody responses. **(a)** Serum neutralizing titers over time were determined by microneutralization (MN) (here shown as mean and SEM of two independent assays). NA inhibition activity was assessed by enzyme linked lectin assay (ELLA) at 4 DPC. **(b)** BAL fluid was taken at 4 DPC and virus neutralization was analyzed by MN, HA specific IgG and IgA titers were measured by ELISA, and NA inhibition was assessed by ELLA. Each data point represents an individual animal. Each serum and BAL sample was assayed twice and a mean computed. **(c)** HA specific memory B cells were detected by ELISpot in PBMC (pre-challenge), spleen and tracheobronchial lymph node (TBLN) 4 DPC. Each animal is represented by a symbol and the mean is shown as a bar. Asterisks denote significance compared to control group (*p<0.05, **p<0.01, *** p<0.001). Serum neutralization was analyzed with two-way ANOVA while Kruskal-Wallis test was used for the analysis of NA neutralisation in sera, BAL samples and ELISpot data.

The neutralizing titer in the BAL was highest in the Aer group (13.4) and neutralizing activity was detectable in 3 of 6 SIM pigs (**Fig. 3b**). There was no detectable neutralization in the BAL of IM immunized animals although haemagglutinin (HA) specific antibodies were present. High levels of anti-HA IgG were present in both IM and SIM groups (119.2 and 167 respectively), while the highest titers of IgA were detected in the Aer and SIM groups (18.3 and 46.6 respectively) (**Fig. 3b**). Only very low levels of NA inhibition were found in BAL compared to serum (**Fig. 3b**). We also evaluated the number of memory IgG and IgA HA specific B cells in spleen, tracheobronchial lymph nodes (TBLN) and blood. Very few antibody secreting cells (ASC) were found in the blood before challenge or in spleen at 4 DPC (**Fig. 3c**). HA specific IgG secreting ASC were detected in TBLN in the Aer (48 ASC/10^6^) and in the SIM (25 ASC/10^6^) groups. A lower number of HA specific IgA ASC were present in these groups. Only 2 of 5 IM immunized pigs had HA specific IgG and IgA ASC in TBLN (**Fig. 3c**).

In summary, IM immunization with S-FLU induced high neutralizing Ab titres in serum, but a limited response in BAL, although HA specific antibodies were present. Aerosol delivery generated the highest neutralizing titers in BAL, but a very low serum response. The SIM group generated a high serum neutralizing titer, although only half the magnitude of IM alone, while the BAL response was lower than in Aer only, but still greater than IM. Statistically significant numbers of HA specific memory B cells were detected only in the Aer group in the local lung lymph nodes.

### Cytokine production by CD4 and CD8 cells in BAL

We analyzed cytokine production of BAL T cells by intracellular staining following *ex vivo* stimulation with H1N1pdm09. No T cell response was detected in the BAL of the IM group. In contrast, Aer and SIM immunization induced a strong T cell response. CD8 T cells in the Aer and SIM groups secreted mainly IFNγ, followed by TNF and the response in both groups was dominated by single IFNγ producers (51%) followed by double secreting IFNγ-TNF (36.2%) and a smaller proportion of triple secreting IFNγ-TNF-IL2 cells (6.5%) (**Fig. 4a**). The only significant CD4 responses were IFNγ in Aer and SIM groups and there were few double or triple cytokine producing cells (**Fig. 4b**). Overall, Aer produced the strongest T cell response, dominated by IFNγ producing cells. SIM induced similar T cells functions although the response was slightly lower in magnitude. Intra-muscular delivery did not generate virus specific T cell in the BAL at 4 DPC.

**Figure 4.**
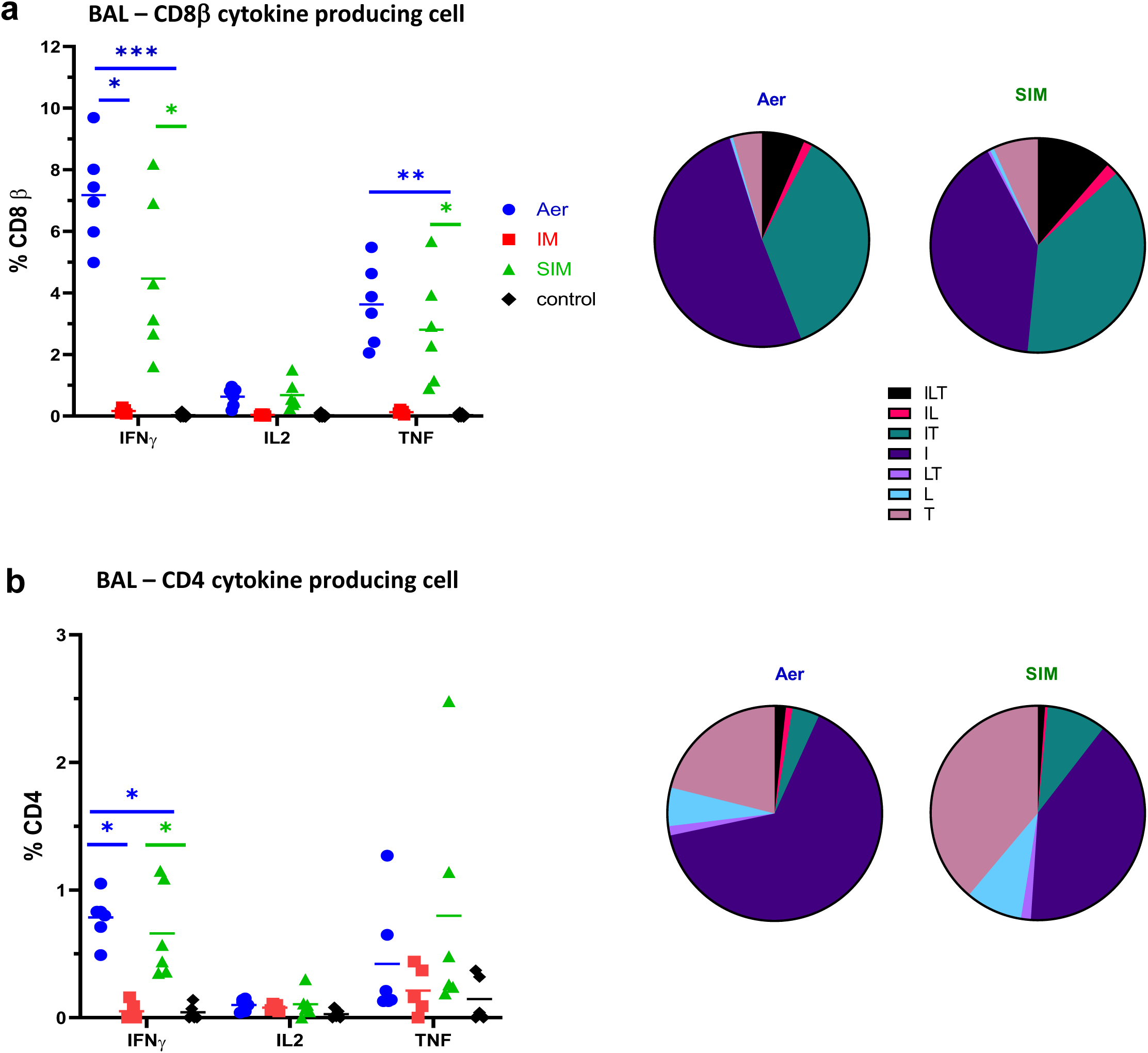
Cytokine secretion in BAL. BAL was collected at 4 DPC. Cryopreserved cells were thawed, stimulated with H1N1pdm09 and IFNγ, IL-2 and TNF cytokine secretion measured in CD8 **(a)** and CD4 **(b)** cells by intracellular staining. Each symbol represent an individual animal and the mean is shown as a bar. The pie chart shows the mean proportion of single, double and triple cytokine secreting CD8 T cells for IFNγ (I), TNF (T) and IL-2 (L). Kruskal-Wallis test was used to compare responses between groups and asterisks indicate significant differences (*p<0.05, **p<0.01, *** p<0.001).

### NP specific tetramer responses in the respiratory tract and blood

We enumerated S-FLU-specific CD8 T cells in blood and different parts of the respiratory tract (nasal turbinates, trachea, BAL and lung) using three NP epitope tetramers: NP_290-298 DFEREGYSL_ (DFE), NP_101-109 NGKWMRELI_ (NGK) and NP_207-225 IAYERMCNI_ (IAY), as previously described^15^ (**Table 1**) (**Fig. 5a**). No responses were detected against the previously identified NP_252-260 EFEDLTFLA_ epitope in all immunized animals (data not shown).

**Table 1.**
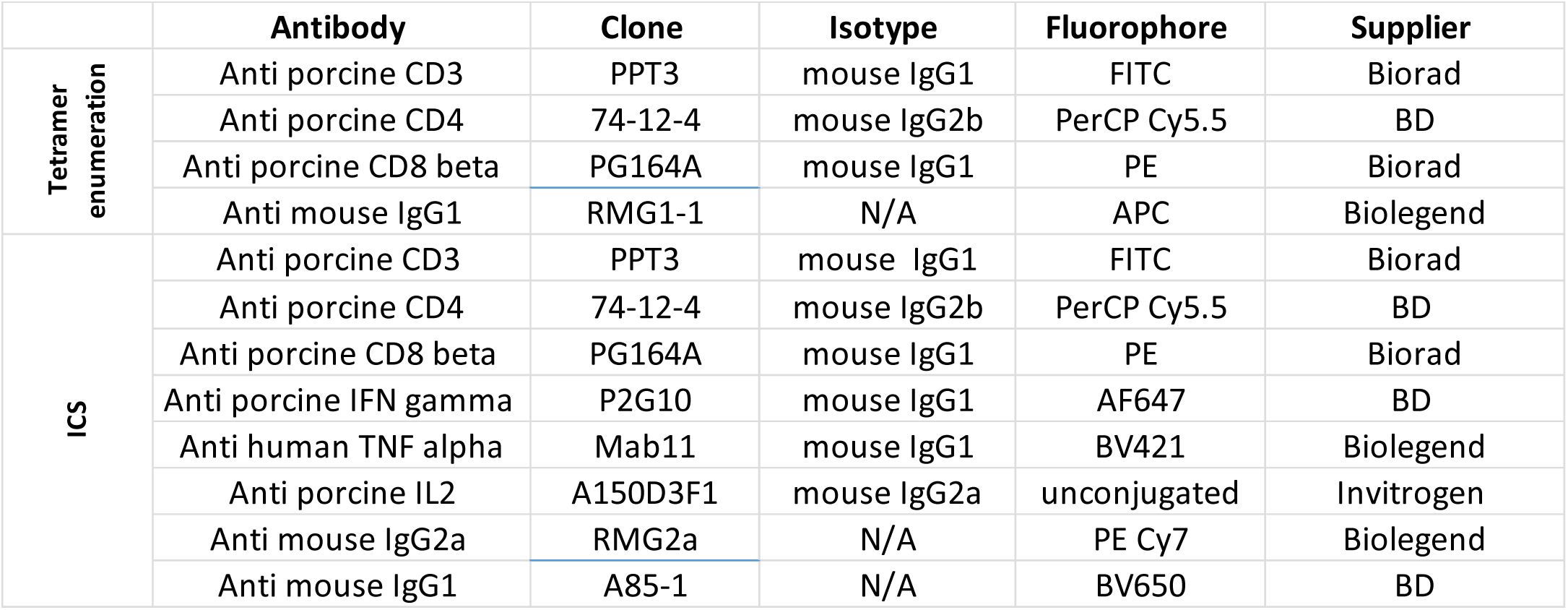
List of antibodies used for intracellular cytokine staining and NP tetramers enumeration.

**Figure 5.**
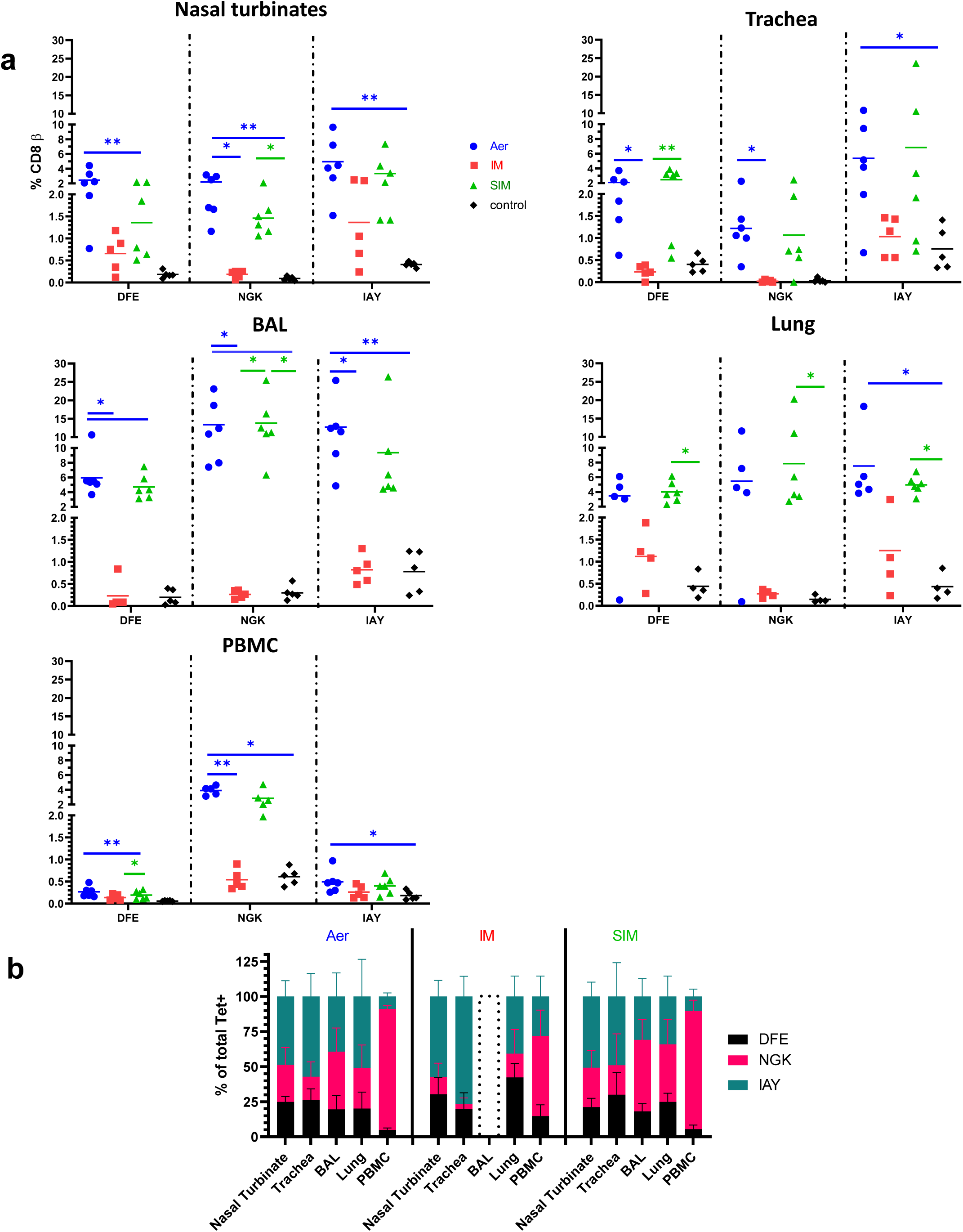
NP specific tetramer responses in respiratory tissues and blood. **(a)** Percentages of DFE, NGK and IAY tetramer + CD8 T cells in the respiratory tract and PBMC. Each symbol represent an individual animal and the mean is shown as a bar **(b)** Proportion of each tetramer among total tetramers+ CD8 T cells in different tissues. A dotted histogram for BAL of the IM group indicates the absence of response. The data represents the average of 2 separate assays. Kruskal-Wallis test was used to compare responses between groups (a) and two-way ANOVA to compare the proportions of tetramers in the different tissues of each group of animals (b). Asterisks denote significant differences (*p<0.05, **p<0.01, *** p<0.001).

In nasal turbinates, the response to IAY was the strongest (5% Aer, 1.4% IM and 3.4% SIM), followed by DFE (2.4% Aer, 0.7% IM, 1.4% SIM) and NGK (2.2% Aer, 0.2% IM, 1.5% SIM). The trachea showed similar specificity. The strongest response was detected in the BAL. NGK^+^ CD8 T cells were the biggest population (13.4% in Aer and 13.8% in SIM) followed by IAY^+^ (12.7% Aer, 9.4% SIM) and a lower response was found to DFE (6% Aer and 4.7% SIM). Strikingly no tetramer staining was found in the BAL of the IM group, in agreement with the lack of intracellular cytokine staining (**Fig. 4**). In the lung similar but lower tetramer specific responses were detected for Aer (5.5% NGK, 7.5% IAY and 3.5% DFE) and the SIM groups (7.9% NGK, 5% IAY and 4.0% DFE) (**Figs. 5a**).

In order to evaluate the hierarchy of tetramer responses in different tissues, we calculated the proportion of each tetramer among total tetramer^+^ CD8 T cells (**Fig. 5b**). The proportions of IAY was much higher in all respiratory tissues compared to blood (p<0.0001 for nasal turbinates compared to PBMC in the Aer group) (**Figs. 5a and b**). The NGK response in blood was greater compared to all respiratory tissues (p<0.0001 when nasal turbinates were compared with blood for the Aer group). In the IM group less NGK^+^ cells were detected in all tissues compared to Aer and SIM. In particular, a significantly lower proportion of NGK^+^ CD8 T cells was found in IM PBMC compared to Aer (p=0.01) although they were still the dominant NP specificity among CD8 T cells (57.1% of total tetramer^+^ CD8 T cells).

Finally, we assessed the numbers of tissue resident memory T cells (TRM) in the respiratory tissues by intravenous infusion of anti-porcine CD3 monoclonal antibody (mAb) as previously described^12^ (**Figs. 6a and b**). The majority of cells in the BAL were inaccessible to the mAb (82.4% average of all 11 animals treated with anti-CD3 mAb) and therefore tissue resident. In the nasal turbinates and trachea 11.6% and 38.8% (average of all 11 animals) of single *ex vivo* labelled CD3 cells (TRM) were detected, while in the lung 95% of the T cells were double positive, perhaps reflecting known difficulties in extracting TRM and blood contamination ^16^. Tetramer positive cells were detected in both TRM and blood borne populations.

**Figure 6.**
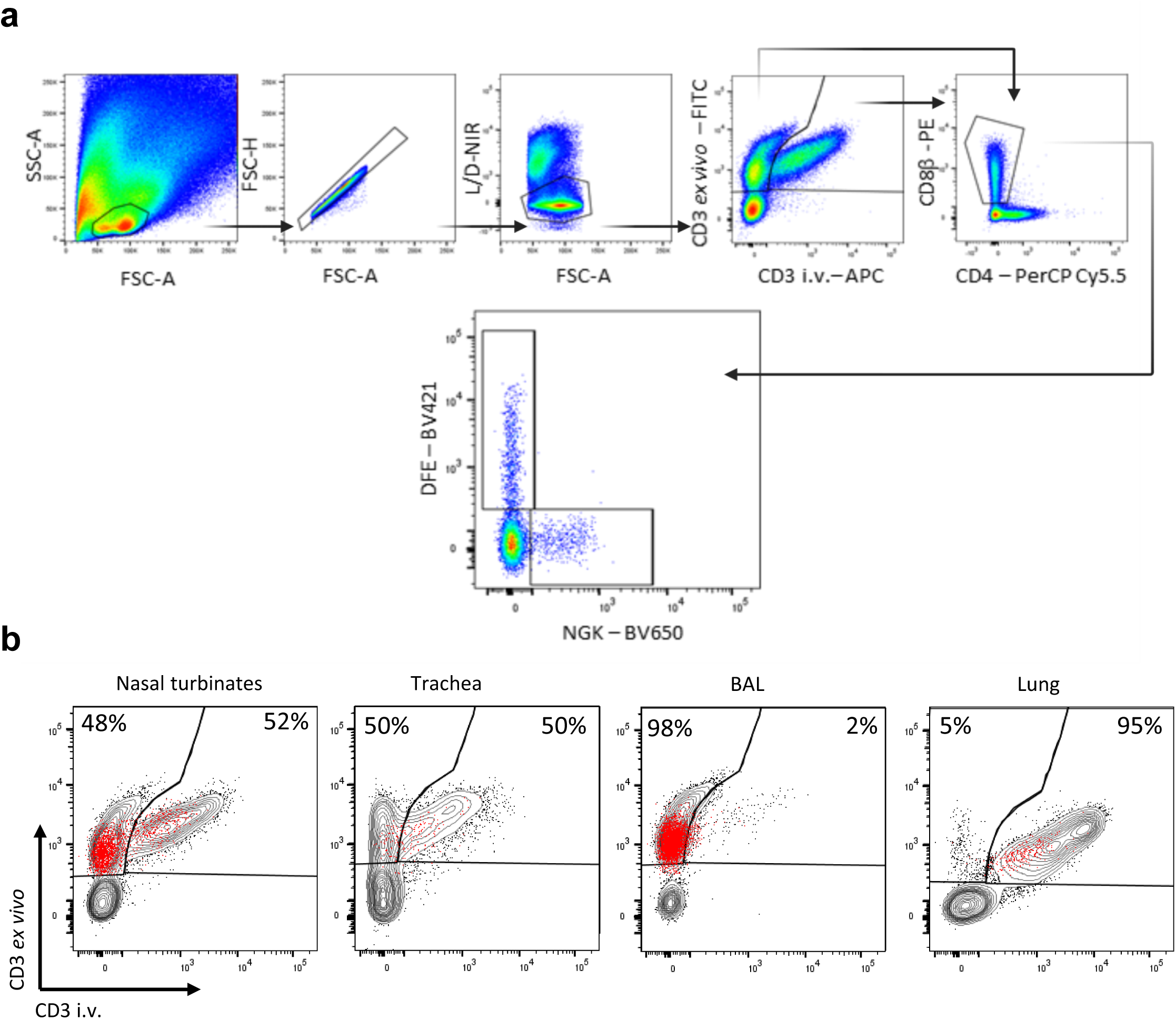
Porcine tissue resident memory cells. Three pigs from the vaccinated groups and two control pigs were infused intra-venously with CD3 mAb and culled 10 min later. **(a)** Representative FACS plots of cells isolated from nasal turbinates of the Aer group showing the gating strategy. **(b)** Lymphocytes were isolated and stained *ex vivo* with the same clone of CD3 Ab labelled with FITC (in grey) as described in the methods. As the infused CD3 does not saturate all CD3 sites some nasal turbinate, tracheal and lung tissue T cells are double positive (intravascular cells). A proportion of BAL, nasal turbinate and tracheal cells are unstained by intravascular mAb, indicating tissue residency. Tetramer positive T cells present after Aer immunization in the different tissues are shown in red.

In summary, we detected strong NP-tetramer specific CD8 T cell responses in the nasal turbinates, trachea, BAL and lung of Aer and SIM immunized animals. IM induced much lower number of tetramer specific cells in all tissues and none in BAL. There was a different hierarchy of the response specificity in the respiratory tract compared to the blood, indicating that sampling blood does not represent responses in the local tissues.

## Discussion

Here we investigated different routes of immunization to determine the most effective in providing protection against influenza A virus in pigs. We tested the traditional systemic (intra-muscular) route used routinely in pigs and humans for influenza immunization and respiratory mucosal immunization, as used with the newer LAIV. We also performed SIM, previously shown to be highly protective, able to induce long lasting immune response and perhaps similar to prime and “pull” immunization regimes that have been investigated recently^5,7,8,17^. In our experiments we administered S-FLU to the whole respiratory tract, a procedure that has been shown in mice to be superior to upper respiratory tract immunization in protecting against heterologous challenge^18^. We were able to do so safely, since S-FLU does not contain a viable RNA segment encoding HA. This obviates the two concerns that dictate restrictions of LAIV to the upper respiratory tract. Firstly, the low level of replication of temperature sensitive LAIV might cause lung pathology and, secondly when used to protect against pandemic influenza viruses reassortment of H5 or H7 with LAIV HA could occur ^9,11,12,19^.

IM immunization induced a powerful neutralizing antibody response, and the viral load and lung pathology were both greatly reduced. However, T cell responses were weak in these animals and strikingly could not be detected in BAL at 4 DPC. Nor were neutralizing antibodies present in BAL, although it should be noted that harvesting BAL involves considerable dilution so low titers of antibodies may be missed. These data indicate that IM immunization fails to develop lung responses, as previously reported in mice with seasonal human inactivated vaccine^20^.

In contrast, in Aer animals, the serum neutralizing titer was much lower but neutralizing activity was detected in BAL. The viral load in nasal swabs was not reduced. There was a trend toward reduced gross and histopathology although this did not reach significance. Aer animals made powerful CD8 and CD4 responses, detectable in BAL, a site containing almost exclusively TRM^12^. Given the powerful T cell and neutralizing antibody responses in the BAL it is surprising that Aer animals showed minimal reduction in nasal virus shedding and a weak effect on pathology, although there was no virus in the BAL. Similarly in a previous experiment pigs immunized by aerosol with H3N2 S-FLU and challenged with heterologous H1N1pdm09 virus exhibited reduced lung pathology 5 DPC, but no reduction in virus shedding ^12^. These data contrast with results in mice where TRM have been shown both to protect against weight loss following heterologous influenza challenge and to reduce viral load^21-23^. This suggests that T cell immunity in mice can both protect against clinical disease (weight loss) and reduce viral load, whereas in pigs a powerful T cells response is insufficient to protect the upper respiratory tract from infection and shedding although the lung viral load and pathology are reduced. Parallel experiments using the same immunization regimes and challenge virus in pigs and ferrets, confirm that small animal models may not always predict the outcome in a large animal natural host, such as the pig ^12^. Correlative studies in humans suggest that cross-reactive T cells provide partial protection against influenza infection ^24-26^. Taken together these data suggest that in large animals, pigs and humans, cross-reactive T cell immunity can ameliorate severe disease but not prevent upper respiratory infection.

The SIM animals showed a combination of the properties of the two immunizations. There was a good serum neutralizing antibody response and neutralizing antibodies were also detected in BAL but, in contrast to IM immunized animals, there was a powerful CD4 and CD8 T cell response in the BAL, lungs and nasal turbinates. SIM animals showed greatly reduced viral load in nasal swabs and no detectable virus in BAL at 4 DPC together with reduced lung pathology. Our study shows that simultaneous systemic and aerosol immunization may have the advantage of providing both protection against homologous challenge, by induction of local and systemic neutralizing antibodies, and heterologous challenge, mediated by local and systemic T cells, including TRM. Systemic immunity may also be of benefit because influenza virus infection can also be systemic and serum neutralizing antibodies may abolish viremia^27,28^. A heterologous challenge of SIM animals would be extremely interesting and confirm whether this immunization regime is truly advantageous.

Simultaneous and prime pull immunization regimes have had mixed success. While systemic prime followed by skin or reproductive tract pull successfully generated protective TRM in these tissues, it has been less easy to protect the respiratory tract by the same strategy^29,30^. Our earlier experiments with BCG in mice and cattle showed that simultaneous parenteral and intra-nasal administration of BCG provided improved protection, which we attributed to earlier local control of mycobacterial replication in the lungs post challenge^5,6^. However, others have not replicated this in primates^31^. Similarly, immunization with an intra-nasal lentiviral Tb vaccine containing antigen 85A (“pull”) after BCG prime failed to improve protection^32^. On the other hand, simultaneous immunization against influenza virus with an adenoviral vector in mice provided improved protection for up to 8 months compared to subcutaneous or intra-nasal immunisation alone and TRM generated by this immunization regime replicated in situ^7,8^. These differing outcomes may be partly explained by the need for additional signals as well as the antigen. For example, an adenoviral but not vesicular stomatitis virus vectored “pull” improved anti-Tb protection in mice^33^ and an adenoviral vector encoding nucleoprotein and 4-1BB ligand enhanced protection against influenza virus, compared to nucleoprotein alone^34^. In the present study, IM S-FLU generated very weak T cell responses and the data provide little evidence for recruitment of systemic T cells to the respiratory tract following infection (“pull”). However, IM S-FLU induce strong neutralizing antibody response, an essential component of influenza vaccines, but the effectiveness of the SIM regime might be improved by a systemic immunization able to generate a stronger circulating memory T cell reservoir, important for the replenishment of local responses ^35^.

While local immune responses are important and, when combined with systemic responses, may provide optimal protection, it is fundamental to know how long they persist. Lung TRM have been shown to be short lived in mice, perhaps as a consequence of the high oxygen tension of the lung microenvironment^35,36^. However, experiments examining the persistence of influenza specific memory indicate that a dividing lung memory population may persist for many months if antigen is retained in the lung^7^. Our data using Ad85A vaccine showed that lung cells from mice immunised 23 weeks previously could stimulate 85A specific T cells to divide, indicating long term persistence of antigen^17^.

Our data illustrate another factor that may partly explain why some regimes have not worked. While we have not analyzed the entire T cell repertoire in detail, the different hierarchy of specificities of tetramer+ CD8 T cells in IM compared to Aer animals suggests that the route of immunization can affect the T cell epitope specificity. Others have found that local cognate antigen recognition is fundamental for establishment of influenza specific TRM ^37,38^ and that local immunodominance is not always found in the circulating T cell pool, although it remains to be shown conclusively whether these differences in immunodominance affect protection^39^.

Our results in this pig influenza challenge model indicate that SIM may offer advantages in protection against influenza viruses. SIM induces an excellent systemic antibody response, known to correlate with protection against homologous virus infection, as well as a powerful local TRM response, vital for protection against heterologous virus challenge. We suggest that development of SIM strategies for other respiratory pathogens including SARS CoV-2, may be advantageous in providing both local protection and a high titer of antibody. SIM strategies should also take into account the need for local co-stimulatory signals and persistence of antigen in the lung.

## Materials and methods

### Vaccine and virus challenge

The H1N1 signal minus influenza vaccine (S-FLU) [eGFP*/N1(A/Eng/195/2009)] H1(A/Eng/195/2009) containing the internal genes of A/Puerto Rico/8/1934 virus was produced as previously described^9^. The swine isolate H1N1 A/swine/England/1353/2009 (H1N1pdm09) was used to infect the pigs.

### Animal immunization and challenge study

The animal experiment was approved by the ethical review process at Animal and Plant Health Agency (APHA) and followed the UK Government Animal (Scientific Procedures) Act 1986. Twenty-four 5-6 weeks old Babraham large white inbred female and male pigs were randomised into four groups of 6 animals as follows: 1) the first group received S-FLU by Aerosol as previously described^12^ (Aer); 2) the second group was immunized intra-muscularly with S-FLU (IM); 3) the third group was immunized simultaneously intra-muscularly and by aerosol with S-FLU (SIM) and 4) unimmunized control group. Two pigs reached their humane end points due to a pre-existing heart condition, limiting the number of pigs in the control and IM groups to 5 animals. During immunization, all the animals were sedated with a cocktail of 4.4mg/kg Zoletil (Virbac, UK) and 0.044mg/kg Domitor (Orion Pharma, UK). Aerosol immunization was performed using a small droplet size vibrating mesh nebuliser (Aerogen Solo, Aerogen Ltd, Ireland) attached to a custom-made veterinary mask^12^. For Aer immunization 2 ml of S-FLU containing 7×10^7^ TCID_50_ S-FLU was administered over 6-10 minutes. For IM administration, the vaccine stock was diluted to a final volume of 4 ml containing 7×10^7^ TCID_50_ and 2 ml were administered to each trapezius muscle behind the ear. Pigs in the SIM group received 2 ml of 3.5×10^7^ TCID_50_ S-FLU by aerosol (as described above) and 3.5×10^7^ TCID_50_ S-FLU delivered in 4 ml IM(2 x 2 ml in each trapezius muscle). The animals were boosted 3 weeks later in a similar manner. Three weeks after the boost, all groups were challenged with 2.8×10^6^ PFU of H1N1pdm09 intranasally using a mucosal atomisation device (MAD300, Wolfe-Tory Medical). For logistic reason, the challenge was performed in 2 different days so that half of the animal in each group were challenged on day 23 post boost and the remaining half on day 25 post boost. Animals were humanely culled at day 4 post challenge (DPC) with an overdose of pentobarbital sodium anaesthetic. At the second cull, 1 mg/kg of anti CD3 purified mAb (PPT3 clone, produced in house) was infused intravenously to the pigs, 10 minutes prior to sacrifice. Since no difference was found in analyses of the samples challenged on different days, the results are presented together. Gross and histopathological analyses were performed as previously described^11,12,40^.

### Tissue Sample Processing

Blood, spleen, broncho-alveolar lavage (BAL) and lung lobes were processed as described previously^11,12^. Trachea and nasal turbinate mucosae were separated from cartilage with tweezers and digested for 2 hours at 37°C in RPMI 1640 supplemented with 100 U/ml penicillin, 100 mg/ml streptomycin, 2 mM L-glutamine (all from Gibco, UK), 2mg/ml collagenase D (Roche, US), 1mg/ml dispase and 1mg/ml of DNase (both from Sigma-Aldrich, UK). Tissues filaments were then mashed with the plunger of a syringe. Isolated cells were then passed through a 70µm cell strainer and red blood cell lysed before cryopreservation in FCS and 10% DMSO. Nasal swabs (one per nostril) were taken daily following infection with H1N1pdm09. Viral titer in nasal swabs and BAL was determined by plaque assay on MDCK cells as previously described^11^.

### Serological assays

ELISA was performed using recombinant HA (from A/England/195/2009) containing a C-terminal thrombin cleavage site, a trimerization sequence, a hexahistidine tag and a BirA recognition sequence as previously described^41^. Microneutralization (MN) was performed using standard procedures as described previously^9,40^.

### Enzyme-Linked Lectin Assay (ELLA)

ELLA was used to quantify neutralization of neuraminidase (NA) enzymatic activity by antibody as described before ^42^. Briefly, NUNC Immuno 96 microwell plates (Sigma-Aldrich, UK) were coated overnight at 4°C with 25 μg/ml fetuin (Sigma-Aldrich, UK) in PBS containing 0.02% sodium azide. Heat inactivated sera and BAL were serially diluted in DMEM supplemented with 0.1% BSA, 100 U/ml penicillin, 100 mg/ml streptomycin and 2 mM L-glutamine starting at 1:40 and 1:4 respectively. H7N1 S-FLU [eGFP/N1(A/Eng/09)] H7(Netherlands/219/2003) was used to minimize any potential steric effect of antibodies binding to H1 HA. An optimal concentration of H7N1 S-FLU was added to the diluted antibodies for 20 minutes on a plate shaker. 100 μl of the mixture of virus and diluted samples were then transferred to the washed coated plate and incubated for 18 hours at 37°C. Peanut agglutinin conjugated with HRP (Sigma-Aldrich, UK) was added at 1 μg/mL in PBS and incubated at room temperature for 2 hours. The plates were washed and developed with 50 µl TMB (Biolegend, UK); after 5 minutes the reaction was stopped with 50 µl 1M sulfuric acid and absorbance measured at 450 and 630 nm. The 50% inhibition titre was calculated as the highest dilution above the 50% inhibition line (midpoint between the signal generated by virus only and medium only wells).

### B cell ELISpot assay

Cryopreserved lymphocytes from blood, spleen and TBLN were used. 10^7^ cells/well were stimulated in each well of a 12 well plate with the TLR7 agonist R484 at 1μg/ml in RPMI 1640 supplemented with 100 U/ml penicillin, 100 mg/ml streptomycin, 10% FBS and 0.1% β mercaptoethanol (all from Gibco, UK). After 48 hours, cells were washed twice with medium and counted. 5×10^5^ cells were distributed in duplicate in assay plates, for the detection of HA specific antibody secreting cells and in negative control wells, while 0.5×10^5^ cells per well were plated to detect all Ig secreting cells (positive controls). Assay plates were MultiScreen™-HA ELISpot plates (Merck, Millipore, UK), coated with anti-porcine IgG, clone MT421 (Mabtech, Sweden), or anti-porcine IgA, A100-102A (Bethyl, US) 1/500 in carbonate buffer overnight at 4°C. After overnight incubation at 37°C, the plates were washed 5 times with PBS containing 0.05% Tween 20 and incubated with biotinylated HA for detection of HA specific B cells (obtained as described before), biotinylated Keyhole limpet hemocyanin (KLH) (Sigma-Aldrich, UK) as a negative control, both at 0.1μg/ml in PBS, or biotinylated anti-porcine IgG (MT424, Mabtech, US) or anti-porcine IgA (A100-102-B, Bethyl, Sweden) at 1/1000 in PBS to detect all Ig secreting cells. After 2 hours incubation, plates were washed and streptavidin alkaline phosphatase (Invitrogen, UK) added for another hour. The plates were then developed and read. Spots detected with KLH were subtracted from the HA response and data presented as antibody secreting cells (ASC) per million cells.

### Flow cytometry

Cryopreserved lymphocytes from BAL were thawed and stimulated with H1N1pdm09 (MOI 1) or medium as a control for 18 hours at 37°C prior GolgiPlug (BD Biosciences, UK) addition as per manufacturer instructions. Following 5 hours incubation with GolgiPlug at 37°C, cells were stained with surface markers (**Table 1**) before fixation and permeabilization using Cytofix Cytoperm (BD Biosciences, UK). Intracellular staining was then performed, and the samples were analysed using an LSRFortessa (BD Biosciences). Data was analysed by Boolean gating using FlowJo v10 (Treestar). For identification of TRM, three animals from each vaccinated group and two control animals were infused i.v. with 1 mg/kg of purified CD3 mAb (clone PPT3) and sacrificed 10 min later, as described above. Cryopreserved lymphocytes isolated from the different tissues were labelled with anti-mouse IgG1-APC, which labels the circulating intravascular cells, for 20 min at 4°C. After two washes with PBS, normal mouse serum was added to block any remaining binding sites of the secondary Ab. The lymphocytes were then stained with surface markers (**Table 1**), including anti-porcine CD3-FITC (clone PPT3, BioRad, UK). As not all CD3 sites would be saturated by intravenous anti-CD3 mAb, circulating T cells are double labelled, while tissue resident T cells are positive only for the *ex vivo* anti CD3-FITC.

NP-tetramer staining was performed on cryopreserved lymphocytes from PBMC, lung, BAL, trachea and nasal turbinate as previously described^15^. Briefly, biotinylated NP peptide loaded SLA monomers were freshly assembled into tetramer with streptavidin BV421 or BV650 (both from Biolegend, UK). Two million mononuclear cells were incubated with protease kinase inhibitor in PBS for 30 minutes at 37°C and tetramers added to the cells on ice for another 30 minutes. Surface staining with optimal antibodies concentration in FACS buffer (PBS supplemented with 2% FCS and 0.05% sodium azide) was performed on ice for 20 minutes (**Table 1**). Samples were washed twice with FACS buffer and fixed in 1% paraformaldehyde before analysis using an LSRFortessa (BD Biosciences).

## Statistical analysis

GraphPad version 8.4.1 was used for statistical analysis. Kruskal-Wallis test was used for the comparison between groups of viral load, pathology, antibody and T cells responses. Two-way ANOVA was used for the comparison of neutralizing antibody and to analyse the hierarchy of the response in the different tissues within the same group.

## Data availability

The datasets generated during the reported study are available on request from the corresponding authors.

## Acknowledgements

We are grateful to the animal staff for excellent animal care. We thank the Pirbright flow cytometry facility for their support and Kelly Roper and Emily Bessell for help with sample processing. We thank APHA for providing the challenge swine A/Sw/Eng/1353/09 influenza virus strain (DEFRA SwIV surveillance programme SW3401).

## Authors contribution

ET, AT, PB, VM conceived, designed and coordinated the study. VM, ET, AM, BP, TC, ME, EM, BC, GD, AS, PB, RM, AT designed and performed experiments, processed samples and analyzed the data. AN carried out postmortem and pathological analysis. ET, PB, VM, AT, wrote the manuscript. All authors read and commented on the manuscript.

